# Assessing microbial growth monitoring methods for challenging strains and cultures

**DOI:** 10.1101/2021.10.05.463247

**Authors:** Damon Brown, Raymond J. Turner

**Author notes:** Corresponding author, (RT).

## Abstract

This is a paper focusing on the comparison of growth curves using field relevant testing methods and moving away from colony counts. Challenges exist to explore antimicrobial growth of fastidious strains, poorly culturable bacterial and bacterial communities of environmental interest. Thus, various approaches have been explored to follow bacteria growth that can be an efficient surrogate for classical optical density or colony forming unit measurements.

Here we tested optical density, ATP assays, DNA concentrations and 16S rRNA qPCR as means to monitor pure culture growth of six different species including *Acetobacterium woodii, Bacillus subtilis, Desulfovibrio vulgaris, Geoalkalibacter subterraneus, Pseudomonas putida* and *Thauera aromatica*. Optical density is and excellent, rapid monitoring method of pure culture planktonic cells but cannot be applied to environmental or complex samples. ATP assays provide rapid results but conversions to cell counts may be misleading for different species. DNA concentration is a very reliable technique which can be used for any sample type and provides genetic materials for downstream applications. qPCR of the 16S rRNA gene is a widely applicable technique for monitoring microbial cell concentrations but is susceptible to variation between replicates. DNA concentrations were found to correlate the best with the other three assays and provides the advantages of rapid extraction, consistency between replicates and potential for downstream analysis, DNA concentrations is determined to be the best universal monitoring method for complex environmental samples.

## Introduction

Assessing growth is fundamental to nearly all microbial studies. On the surface this is an easy procedure done in introductory courses worldwide [1]. However, it turns out this routine experiment is not as trivial as one thinks. For easily culturable aerobic species, the process is relatively simple as the growth medium need only contain the appropriate carbon sources and essential nutrients to culture the typical well studied model microbes. After the appropriate growth medium has been selected, direct cell counting on agar plates can be performed for accurate quantification providing colony forming units (CFU) or viable cell count (VCC) values. However, it has become apparent that this method restricts the scope of species possible for study and cannot but use to study complex environments [2–4]. For more rapid analysis, optical density (OD) is evaluating the scatting of light by cells, either using the classical Klett meter or an absorption spectrometer set to 550 or 600 nm. This however is also limiting due to suspended material in various growth media and the inability to decipher living vs dead vs flocculation from extracellular polysaccharides. Additionally, size and shape of cells effect scattering and can lead to misinterpretation of cell numbers. Related rapid assays includes color dependant activity assays (colloquially known as bug bottles or biological activity reaction test (BART) bottles [5]), microscopy (grid cell counting and live dead staining) [6–8], dyes such as crystal violet or 5-(4,6-dichlorotriazinyl) aminofluorescein (DTAF) for total biomass staining [9,10] or metabolic dyes to determine actively respiring cells such as 5-cyano-2,3-ditolyl tetrazolium chloride (CTC) [10]. Fluorescent *in situ* hybridization (FISH) is a target-specific approach which relies on a fluorescent reporter attached to a nucleic probe to determine the presence and abundance of the target sequence. This can be used for total or genera-specific cell enumeration when targeting a gene such as 16S rRNA [11,12]. Now in our genomics era, 16S rRNA quantification using quantitative PCR (qPCR) is gaining popularity with [13] or without sequencing databases [14,15].

For anaerobes, accurate cell enumeration becomes increasingly more difficult as different anaerobes will require different oxidation-reduction potentials (ORP) to thrive; ≤ −100 mV for obligate anaerobes [16] and ≤ −330 mV for strict anaerobes [17]. Following the improved Hungate culturing technique of anaerobes [18,19], it is possible to isolate and enumerate pure culture anaerobes [20,21] but direct microscopy and fluorescent in-situ hybridization (FISH) and more common enumeration techniques [22–25]. A review of anaerobic culturing and quantification is available elsewhere [26]. In many instances, the difficulty of culturing anaerobes forces researchers to choose indirect methods to assess growth and activity, such as rates of substrate consumption or end-product production, where an easily quantified chemical is sampled and measured at time points in favor of actual cell counting [27,28]. The rate of consumption or production relates to cell growth through the Monod equation [29]. It should be noted that there is a distinction between cell counting and microbial activity, and the need to monitor one or the other or both depends on the research, environmental, industrial, or medical question being asked.

The above exemplify the issues of accurate quantification of aerobic and anaerobic pure cultures. This task becomes exponentially more difficult when considering environmental samples which have diverse, unspecified microbial species present. In such examples, highly specific monitoring methods are no longer viable to determine accurate cell counts so general techniques must be used. The typical trade off when transitioning to general monitoring from selective is the loss of specificity (e.g. the presence of a specific pathogen) for the gain of total cell counts. General quantification is an important technique for the monitoring of hydrocarbon remediation efforts and wastewater treatments [30,31].

In this study, we take the opportunity to review the various methods to evaluate microbial growth. We then compare select testing methods used in environmental/industrial applications. From our groups interest we will look at ATP levels and 16S rRNA qPCR which are being adapted in in oil and gas industries in western Canada. These are then compared to traditional OD_600_ and qPCR targeting 16S rRNA to determine how reliable, complementary and efficient these differing methods are.

## Materials and methods

### Cultures and media

Six pure cultures were used in this study acquired from DSMZ. The cultures are *Acetobacterium woodii* (DSM 1030), *Bacillus subtilis* (DSM 10), *Desulfovibrio vulgaris* (DSM 644), *Geoalkalibacter subterraneus* (DSM 29995), *Pseuodomonas putida* (DSM 291), and *Thauera aromatica* (DSM 6984). Each of the chosen species has a representative genome sequenced on NCBI and their details are in Table 1. Pure cultures were recovered from −70 °C freezer stocks (10% glycerol) into the suggested media (DSM medium 135, medium 1, medium 63, medium 1249, medium 1a, and medium 586, respectively) prepared in 20 mL aliquots and sealed in 26 mL Hungate tubes with anaerobic headspaces (either N_2_ or N_2_/CO_2_). Fresh media tubes were inoculated with the freezer recovery culture and incubated at the recommended temperatures for 7 days, then fresh media tubes were inoculated in triplicate with a 10% inoculant.

**Table 1.** Representative sequenced genome chosen and identified 16S rRNA copy numbers.

| Species and strain | NCBI accession number | Genome length (bp) | 16S rRNA copy number* |
| --- | --- | --- | --- |
| *Acetobacterium woodii* DSM 1030 | CP002987.1 | 4044777 | 5 |
| *Bacillus subtilis subtilis* 168 | CP010052 | 4215619 | 10 |
| *Desulfovibrio vulgaris* Miyazaki F | CP001197.1 | 4040304 | 4 |
| *Geoalkalibacter subterraneus* Red1 | CP010311 | 3475523 | 4 |
| *Pseudomonas putida* KT2440 | AE015451.2 | 6181873 | 7 |
| *Thauera aromatica* MZ1T | CP001281.2 | 4496212 | 4 |

\*Counted manually from NCBI sequences

### Sampling and testing

Once inoculated, the fresh cultures were sampled in a time course for testing. At each point, sterile syringe and needles were used to aseptically remove 2 mL. This aliquot was split, 1 mL used to measure OD_600_ in a Hitachi U-2000 Spectrophotometer (reference sample was uninoculated media) and was then recovered and used for DNA extraction in the MPBio FastDNA^®^ Spin Kit. The DNA concentration was measured using Invitrogen Qubit™ fluorometer and the Quant-iT™ dsDNA HS assay kit. Following quantification, DNA was cleaned using the OneStep™ PCR inhibitor removal kit (Zymo Research) prior to being used in quantitative PCR (qPCR). The other 1 mL was consumed to measure ATP using the LifeCheck ATP test kits (OSP).

### Quantitative PCR

qPCR was done targeting the 16S rRNA gene, specifically 515-809 (variable region 4) using modified primers from Caporaso *et al.* [32]. Primer sequences are provided in Table 2. Starting quantification was determined using synthetic gBlocks purchased from IDT (IDTdna.com) at concentrations of 10^8-3^ copies/μL. The gBlock used contained the target 16S rRNA sequence flanked by two multidrug resistance genes (for use in other studies), separated by sequences of ten thymines. The entire gBlock sequence is provided in Table 3. Reaction mixtures were prepared to a total volume of 20 μL, with 10 μL PowerUp™ SYBR™ Green 2X Master Mix (appliedbiosystems), 1.2 μL 10 μM 515_F, 0.6 μL 10 μM 806_R, 6.2 μL nuclease free water and 2 μL DNA template. Thermocycling was performed in a BioRad CFX96 Real-Time PCR System with the following protocol: 50 °C – 2 minutes, 95 °C – 2 minutes followed by 50 cycles of 95 °C for 45 seconds, 55 °C for 30 seconds and 72 °C for 45 seconds.

**Table 2.** Primer sequences used in quantitative PCR.

| Primer name | Sequence (5'-3') | Melting Temperature (°C) |
| --- | --- | --- |
| 519_F | CAG CMG CCG CGG TAA | 57.6 |
| 806_R | GGA CTA CHV GGG TWT CTA AT | 50.7 |

Nucleotide codes: H = A/C/T; M = A/C; V = A/C/G; W = A/T

**Table 3.**
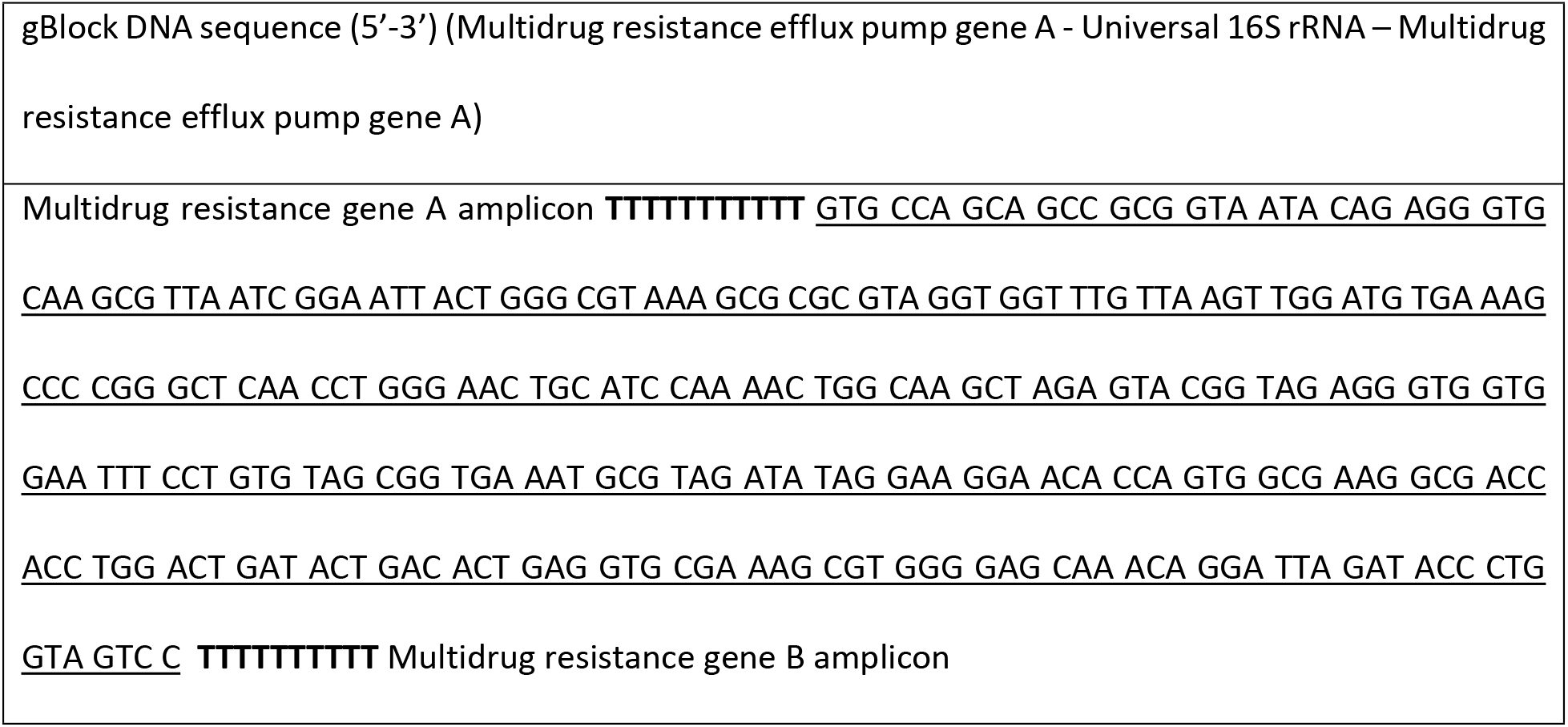
gBlock DNA sequence.

### Cell count calculations

For direct comparison, cell counts, such as they are inferred and calculated for each method, were calculated. OD_600_ was not converted to cell count equivalent in presented data. The chosen ATP method produces relative light units (RLU) per milliliter, as 1 mL of each culture was used in the assay, the microbial equivalents (ME) produced are used directly as calculated cell counts. DNA concentrations were converted into a cell count proxy using the assumption of 2 fg of DNA per cell, taken as an average from values reported by Bakken and Olson, 1989 [33]. To convert qPCR values into cell counts, the copies of 16S rRNA genes per μL were converted to copies mL^−1^, then divided by the 16S rRNA gene copies counted in the NCBI sequenced genomes (see Table 1).

## Results

### Optical Density

Growth of each culture was monitored using OD_600_ (reference media was each species’ uninoculated media). The results of the time course sampling using OD_600_ for each species is shown in Figure 1, panels A-F. It is important to note the time courses and the OD_600_ scales are all unique for each species. *A. woodii, G. subterraneus* and *P. putida* (Fig. 1A, D, and E) followed a typical sigmoidal growth curve although their lag phases varied in length. *B. subtilis* (Fig. 1B) showed a diauxic growth curve, likely a result of the media not being fully anoxic but containing an anaerobic headspace. *D. vulgaris* is a steady increase over the course of the 48-hour testing period rather than a sharp sigmoidal curve (Fig. 1C). Is it interesting to note that the growth trend is still observable, though muted, in the *D. vulgaris* culture despite having readings above 1.0 (Figure 1.C). *T. aromatica* (Fig. 1F) shows a modified sigmoidal curve, however the peak occurs following the log phase before dipping then and plateauing into the stationary phase.

**Figure 1.**
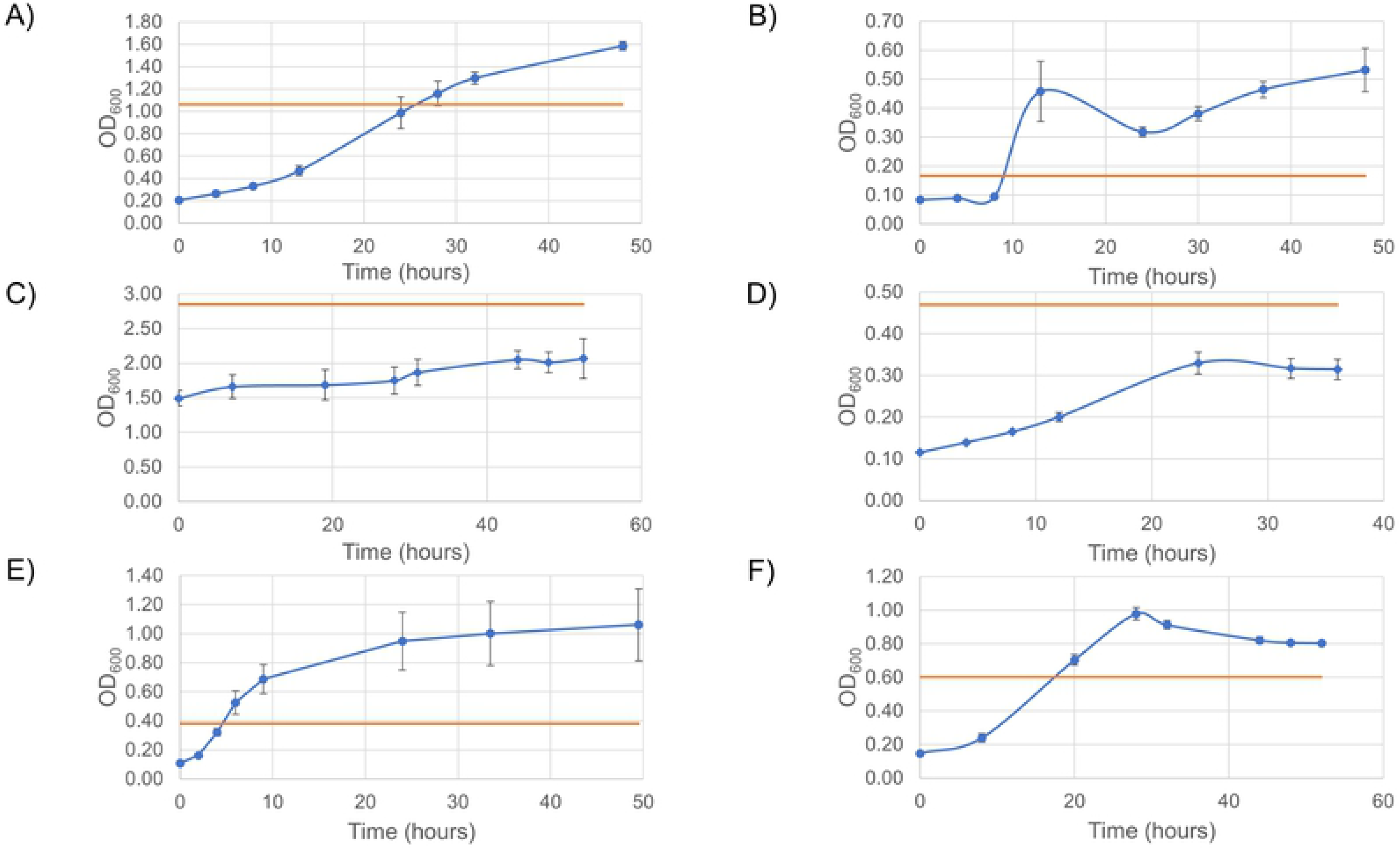
Optical density time course for each species. OD_600_ time course of A) *A. woodii,* B) *B. subtilis,* C) *D. vulgaris,* D) *G. subterraneus,* E) *P. putida* and F) *T. aromatica*. Blue lines are the species growth curves (n=3), orange line is the inoculant value.

A formula was created by Kim et al. (2012) studying *Pseudomonas aeruginosa* which found the following relation of OD_600_ to colony forming units (CFU):

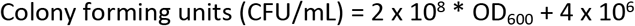

It is noted that this formula should be confirmed by plating cells at unique OD_600_ values and validating the formula for each pure culture. It is acknowledged that the differences in culture turbidity of the six species used in this study and the inability to grow all six on plated media means converting OD_600_ to CFU values will be an inaccurate conversion, but this was still done to maintain uniformity between datasets and to highlight the issue of using such conversion factors without confirming and modifying the equation empirically for each species. Thus we used this to convert values for time zero, a time point to represent the mid log phase and a time point to represent the stationary phase, which are reported in Table 5.

**Table 5.** Summary of the calculated cell counts per milliliter from each monitoring method at the initial time point, mid log phase and stationary phase for all species tested.

| Species | Time point<br>(hours) | Testing method |  |  |  |
| --- | --- | --- | --- | --- | --- |
|  |  | Converted<br>CFU/mL from<br>OD<sub>600</sub> | ATP (ME/mL) | Calculated cells<br>from DNA<br>(cells/mL) | Calculated<br>cells from 16S<br>rRNA<br>(cells/mL) |
| *A. woodii* | 0 | $4.55 \times 10^7 \pm$<br>$6.60 \times 10^5$ | $8.77 \times 10^7 \pm$<br>$8.29 \times 10^6$ | $3.02 \times 10^8 \pm$<br>$2.16 \times 10^7$ | $1.45 \times 10^8 \pm$<br>$3.48 \times 10^7$ |
| | 13 | $9.77 \times 10^7 \pm$<br>$8.77 \times 10^6$ | $2.74 \times 10^8 \pm$<br>$7.05 \times 10^7$ | $3.99 \times 10^9 \pm$<br>$6.71 \times 10^8$ | $1.42 \times 10^9 \pm$<br>$1.96 \times 10^8$ |
| | 32 | $2.63 \times 10^8 \pm$<br>$1.12 \times 10^7$ | $1.45 \times 10^9 \pm$<br>$3.74 \times 10^8$ | $1.18 \times 10^{10} \pm$<br>$1.35 \times 10^9$ | $3.41 \times 10^9 \pm$<br>$2.92 \times 10^8$ |
| *B. subtilis* | 0 | $2.07 \times 10^7 \pm$<br>$4.99 \times 10^5$ | $3.23 \times 10^4 \pm$<br>$2.98 \times 10^4$ | $2.71 \times 10^8 \pm$<br>$6.29 \times 10^7$ | $8.85 \times 10^7 \pm$<br>$1.79 \times 10^7$ |
| | 13 | $9.57 \times 10^7 \pm$<br>$2.08 \times 10^7$ | $2.64 \times 10^8 \pm$<br>$1.08 \times 10^8$ | $3.28 \times 10^9 \pm$<br>$1.79 \times 10^9$ | $8.44 \times 10^8 \pm 3.79$<br>$\times 10^8$ |
| | 30 | $8.03 \times 10^7 \pm 4.76 \times 10^6$ | $2.30 \times 10^8 \pm 5.61 \times 10^7$ | $4.98 \times 10^9 \pm 2.40 \times 10^9$ | $8.40 \times 10^8 \pm 4.24 \times 10^8$ |
| *D. vulgaris* | 0 | $3.03 \times 10^8 \pm 2.26 \times 10^7$ | $5.59 \times 10^6 \pm 9.61 \times 10^4$ | $7.32 \times 10^9 \pm 8.65 \times 10^9$ | $7.24 \times 10^8 \pm 3.32 \times 10^8$ |
| | 28 | $3.54 \times 10^8 \pm 3.88 \times 10^7$ | $2.00 \times 10^8 \pm 4.28 \times 10^7$ | $1.08 \times 10^9 \pm 2.41 \times 10^8$ | $4.08 \times 10^9 \pm 8.04 \times 10^8$ |
| | 48 | $4.06 \times 10^8 \pm 2.96 \times 10^7$ | $1.53 \times 10^8 \pm 5.43 \times 10^7$ | $1.78 \times 10^{10} \pm 1.10 \times 10^9$ | $9.08 \times 10^9 \pm 1.93 \times 10^9$ |
| *G. subterraneus* | 0 | $2.71 \times 10^7 \pm 9.43 \times 10^4$ | $8.15 \times 10^7 \pm 8.02 \times 10^6$ | $2.23 \times 10^9 \pm 2.24 \times 10^8$ | $3.77 \times 10^9 \pm 4.51 \times 10^8$ |
| | 12 | $4.41 \times 10^7 \pm 2.19 \times 10^6$ | $2.41 \times 10^8 \pm 8.28 \times 10^7$ | $6.00 \times 10^9 \pm 7.82 \times 10^8$ | $6.12 \times 10^9 \pm 6.02 \times 10^8$ |
| | 32 | $6.74 \times 10^7 \pm 4.81 \times 10^6$ | $9.16 \times 10^7 \pm 1.75 \times 10^7$ | $1.43 \times 10^{10} \pm 2.95 \times 10^9$ | $1.17 \times 10^{10} \pm 2.62 \times 10^9$ |
| *P. putida* | 0 | $2.58 \times 10^7 \pm 1.63 \times 10^5$ | $4.12 \times 10^7 \pm 2.55 \times 10^6$ | $2.09 \times 10^9 \pm 1.21 \times 10^8$ | $4.29 \times 10^8 \pm 1.47 \times 10^8$ |
| | 6 | $1.09 \times 10^8 \pm 1.61 \times 10^7$ | $5.77 \times 10^8 \pm 1.08 \times 10^8$ | $2.47 \times 10^{10} \pm 5.26 \times 10^9$ | $7.96 \times 10^9 \pm 2.59 \times 10^9$ |
| | 33.5 | $2.04 \times 10^8 \pm 4.39 \times 10^7$ | $8.65 \times 10^8 \pm 1.46 \times 10^8$ | $3.99 \times 10^{10} \pm 9.48 \times 10^9$ | $4.79 \times 10^9 \pm 1.55 \times 10^9$ |
| *T. aromatica* | 0 | $3.35 \times 10^7 \pm 1.39 \times 10^6$ | $5.81 \times 10^7 \pm 2.97 \times 10^6$ | $2.90 \times 10^{10} \pm 4.08 \times 10^8$ | $7.01 \times 10^8 \pm 2.83 \times 10^8$ |
| | 20 | $1.45 \times 10^8 \pm$<br>$6.38 \times 10^6$ | $3.51 \times 10^8 \pm$<br>$2.64 \times 10^7$ | $6.11 \times 10^8 \pm$<br>$1.11 \times 10^8$ | $1.41 \times 10^{10} \pm$<br>$7.54 \times 10^9$ |
| | 48 | $1.65 \times 10^8 \pm$<br>$2.30 \times 10^6$ | $4.41 \times 10^8 \pm$<br>$2.90 \times 10^7$ | $1.41 \times 10^{10} \pm$<br>$1.53 \times 10^9$ | $2.62 \times 10^{10} \pm$<br>$7.66 \times 10^9$ |

### Cellular ATP Levels

ATP measurements were converted into microbial equivalents (ME) according to the manufacturer’s protocol using the relative light units (RLU) from manufacturer’s luminometer as in equation (1).

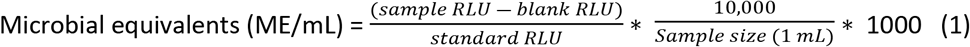

Calculated ME are plotted in Figure 2. Data is not displayed in log scale to better illustrate fluctuations in data trends. *A. woodii* shows a gradual increase in ME/mL between time 0 and 24 hours, going from 8.77 × 10^7^ ME/mL (T = 0) to 7.48 × 10^8^ ME/mL at 24 hours (Fig. 2A). There was a sharp increase after this point to 1.45 × 10^9^ ME/mL at 32 hours, after which readings remained relatively stable at 1.33 × 10^9^ ME/mL until the final time point of 48 hours. *B. subtilis* readings were less constant, reaching peak values at 13 hours (2.64 × 10^8^ ME/mL) before dipping to 8.26 × 10^7^ ME/mL at 24 hours and subsequently recovering to 2.47 × 10^8^ ME/mL at 37 hours. After this point readings gradually decrease to 1.58 × 10^8^ ME/mL at the final timepoint of 48 hours (Fig. 2B). *D. vulgaris* ME/mL readings did not follow a typical sigmoidal growth curve, rather they peaked at 31 hours (3.01 × 10^8^ ME/mL) before declining to 1.56 × 10^8^ ME/mL at 44 hours where it remained relatively stable for the remainder of the time points (Fig. 2C). *G. subterraneus* had a similar trend in ME, with the peak occurring at 12 hours (2.41 × 10^8^ ME/mL) before decreasing to 6.30 × 10^7^ ME/mL at the final time point (T = 36 hours) (Fig. 2D). *P. putida* followed a sigmoidal curve with a short lag phase in the first two hours (4.12 × 10^7^ ME/mL at T = 0 to 1.17 × 10^8^ at T = 2 hours) before reaching 5.77 × 10^8^ ME/mL at 6 hours, after which it gradually increased for the remainder of the growth curve, reaching 9.89 × 10^8^ ME/mL at T = 49.5 hours (Fig. 2E). *T. aromatica* followed a sigmoidal curve with a lag phase between 0 and 8 hours (5.81 × 10^7^ ME/mL and 9.15 × 10^7^ ME/mL, respectively) before increasing to 6.29 × 10^8^ ME/mL at T = 32 hours where it remains until T = 44 hours, after which it drops to 4.41 × 10^8^ ME/mL at T = 48 hours (Fig. 2E).

**Figure 2.**
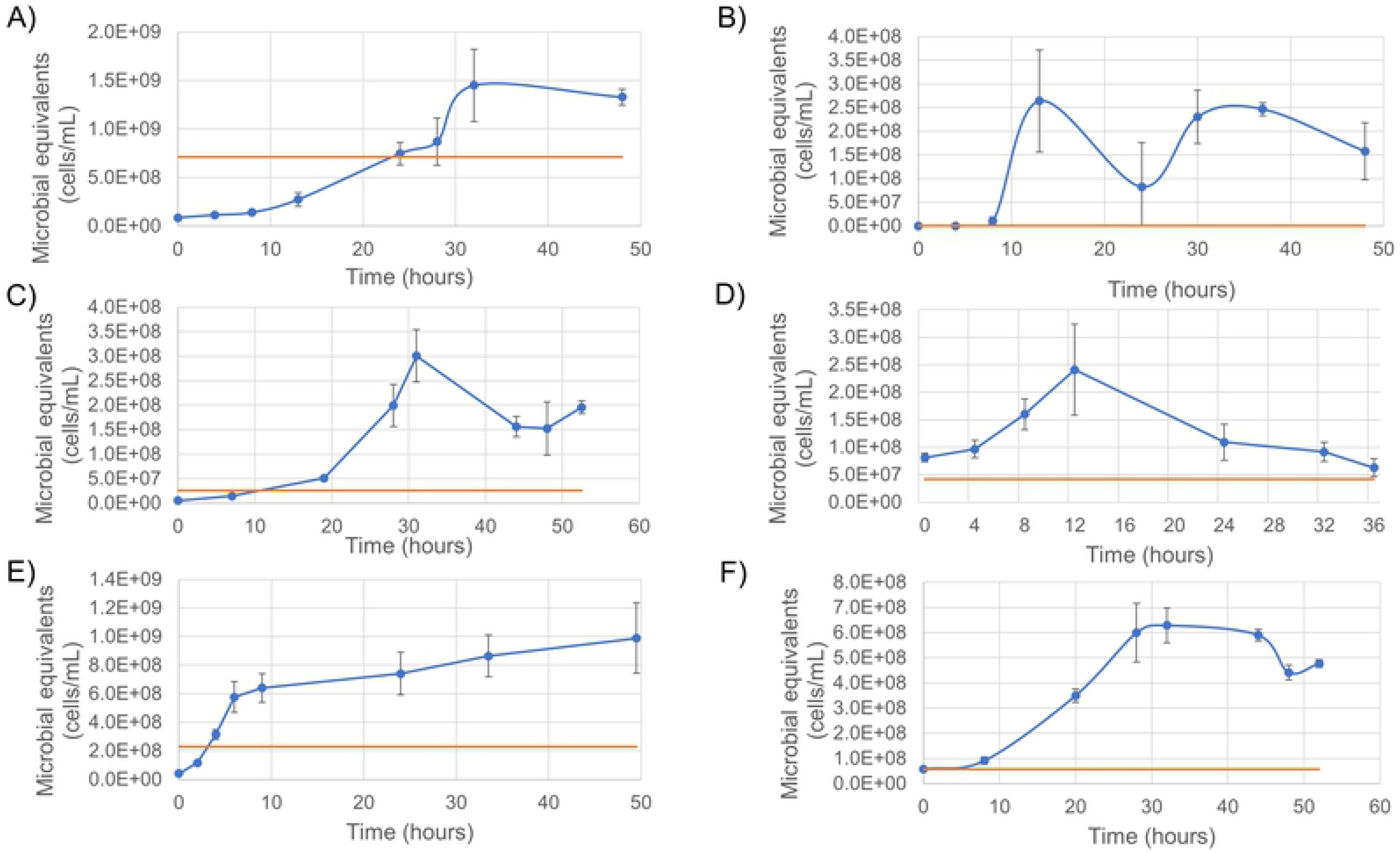
Microbial equivalents time course as determined using luciferase-based ATP assay for each species. Microbial equivalents per milliliter calculated for a time course of A) *A. woodii,* B) *B. subtilis,* C) *D. vulgaris,* D) *G. subterraneus,* E) *P. putida* and F) *T. aromatica*. Blue lines are the species growth curves (n=3), orange line is the inoculant value.

An interesting observation is the ATP values of the T = 0 hour time point is higher or near to the value of the inoculant, as in the case of *G. subterraneus* (Fig. 2D), *T. aromatica* (Fig. 2F) and *P. putida* (Fig. 2E). It is hypothesized this is a result of the inoculant culture being in stationary and/or death phase at the time of inoculation, lowering the ATP readings, but the cultures were capable of rapidly activating their metabolism again when exposed to fresh media. This highlights that ATP levels reflect metabolizing bacteria concentration only.

### DNA concentration

The DNA concentrations were measured from 1 mL aliquots sampled at each time point and are reported in Figure 3. *A. woodii* DNA concentrations follow a typical sigmoidal curve but peaked at the mid-point of the growth curve (24.9 μg/mL, T = 24 hours), after which the DNA concentration dipped slightly to 21.1 μg/mL at T= 28 hours then remained stable between 23.6 μg/mL and 27.5 μg/mL (Fig. 3A). *B. subtilis* DNA concentrations followed a similar trend as the OD_600_ and ATP, indicating a diauxic growth pattern (Fig. 3B), where DNA concentrations were at their highest at 24 and 48 hours (11.8 μg/mL and 12.0 μg/mL respectively). Between these time points the DNA concentrations dip to 7.2 μg/mL at 37 hours. *D. vulgaris* DNA concentrations followed more of a sigmoidal curve compared to the OD_600_ readings, possibly owing to the high scattering properties of the *D. vulgaris* media particulates minimizing the effect of the cells alone, as discussed above. DNA concentrations showed a lag phase between T = 0 hours and T = 19 hours, where concentrations varied between 1.7-2.7 μg/mL before increasing to 28.7 μg/mL at T = 31 hours (Fig. 3C). DNA concentrations peaked at 44 hours (37.0 μg/mL) before decreasing to 30.0 μg/mL at the final time point (T = 52.5 hours). *G. subterraneus* DNA concentrations followed a standard sigmoidal curve, nearly identical to the OD_600_ curve. The lag phase occurred between 0 and 8 hours, during which DNA concentrations were between 4.2-6.1 μg/mL before increasing up to 32.7 μg/mL at 24 hours and decreasing to 28.5 μg/mL at 32 hours (Fig. 3D). It should be noted DNA was not collected at the 36 hour time point due to lack of supplies, thus DNA concentrations could not be collected for the final time point. *P. putida* also exhibited a classic sigmoidal curve in the DNA concentrations as with the OD_600_ and ATP, but there is a shorter lag phase and a longer stationary phase (Fig. 3E) than is typical of a sigmoidal curve. The lag phase was between 0 and 2 hours (4.2 μg/mL and 8.9 μg/mL, respectively), then DNA concentrations increased to 63.9 μg/mL at 9 hours, then gradually increasing to 86.4 μg/mL by the final time point at 49.5 hours. *T. aromatica* DNA concentrations follow a gradual sigmoidal curve, lacking significant lag or stationary phases (Fig. 3F). The lag phase was between 0 and 8 hours (1.2 μg/mL and 3.2 μg/mL respectively), then a steady increase to 57.0 μg/mL by 48 hours, after which there was a slight decline to 55.3 μg/mL at the last time point.

**Figure 3.**
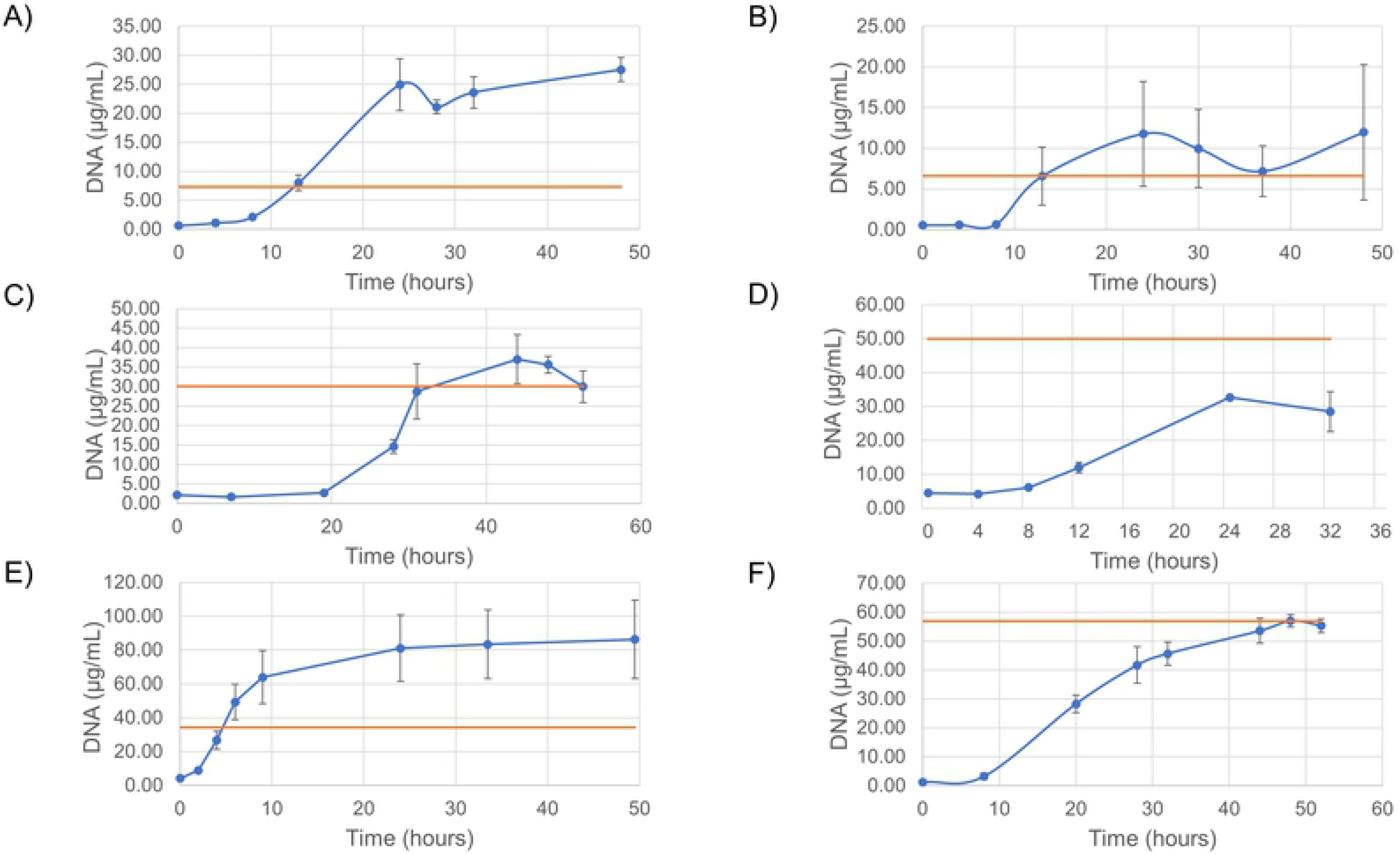
DNA concentration time course for each species. DNA concentrations in micrograms per milliliter over a time course of A) *A. woodii,* B) *B. subtilis,* C) *D. vulgaris,* D) *G. subterraneus,* E) *P. putida* and F) *T. aromatica* as measured by a Qubit fluorometer. Blue lines are the species growth curves (n=3), orange line is the inoculant value.

DNA concentrations were converted into a cell proxy estimation by assuming 2.0 fg DNA per cell based on the average 1.6-2.4 fg/cell [33]. Converted cell proxy values for time zero, a time point representative of mid log phase and a time point representative of the stationary phase are reported in Table 5. An interesting point is that at all time points, even initial (0 hours), the calculated cell counts are never below 10^8^ cells/mL, and between initial time points and stationary phase, the calculated cell counts increase by one or two orders of magnitude.

### qPCR of 16S rRNA

As with the ATP data, we have not plotted the qPCR data in log format. Each of the three biological replicates collected at each timepoint were ran in duplicate on the thermocycler, raising the N value to six. The 16S rRNA copies measured per μL are shown in Figure 4. A key point to highlight in all the trends is an increase in the error bars, especially in the later time points of the graphs. The trends of the 16S rRNA qPCR results followed the DNA concentrations for all species. The 16S rRNA copy numbers for *A. woodii* showed the same lag phase as DNA between 0 and 8 hours, where copy numbers were between 7.23 × 10^5^ copies/μL and 1.69 × 10^6^ copies/μL. The counts increase to 1.64 × 10^7^ copies/μL at 24 hours, then dip slightly to 1.58 × 10^7^ copies/μL before increasing again to 2.07 × 10^7^ copies/μL at the final time point of 48 hours (Fig. 4A). *B. subtilis* showed a lag phase up to 8 hours (3.09 × 10^5^ – 8.85 × 10^5^ copies/μL) before falling into the same diauxic growth curve seen in other monitoring methods. The initial increase peaked at 24 hours (1.11 × 10^7^ copies/μL) before falling to 6.04 × 10^6^ copies/μL at 37 hours and increasing again to 1.13 × 10^7^ copies/μL at 48 hours (Fig. 4B). The lag phase of *D. vulgaris* occurred between 0 and 19 hours (2.35 × 10^6^ – 4.04 × 10^6^ copies/μL) before increasing to 3.99 × 10^7^ copies/μL at 31 hours, where they remained relatively stable until 48 hours (3.63 × 10^7^ copies/μL) (Fig. 4C). At this point, the *D. vulgaris* curve becomes skewed due to significant error bars on the final time point (T = 52.5 hours), owing to half the replicates (n = 3) reporting values an order of magnitude less than the other three replicates (1.52 × 10^8^ copies/μL ± 1.16 × 10^8^). The *G. subterraneus* growth curve shows a lag phase between 0 and 8 hours (1.17 × 10^7^ – 1.72 × 10^7^ copies/μL) before increasing to 5.38 × 10^7^ copies/μL at 24 hours then decreasing to 4.70 × 10^7^ copies/μL at 32 hours (Fig. 4D). As with the DNA concentrations, no DNA was available for the final time point (T = 36 hours) and therefore no values are reported. *P. putida* 16S rRNA copies/μL show a far less tidy sigmoidal curve compared to the other monitoring methods. The lag phase occurred between 0 and 2 hours (3.00 × 10^6^ – 9.64 × 10^6^ copies/μL) before it increased to 5.57 × 10^7^ copies/μL at the start of stationary phase (T = 6 hours) (Fig. 4E). After T = 6 hours where other lines of evidence show a gradual increase into a plateau, the 16S rRNA data fluctuate between 3.36 × 10^7^ – 7.20 × 10^7^ copies/μL with the peak values occurring at T= 24 hours (7.20 × 10^7^ copies/μL). *T. aromatica* 16S rRNA copies/μL show a lag phase between 0 and 8 hours (2.80 × 10^6^ – 6.79 × 10^6^ copies/μL) before increasing to 9.57 × 10^7^ copies/μL at 28 hours then enter stationary phase for the remainder of the growth curve, which increased slightly to 1.05 × 10^8^ copies/μL at 48 hours (Fig. 4F).

**Figure 4.**
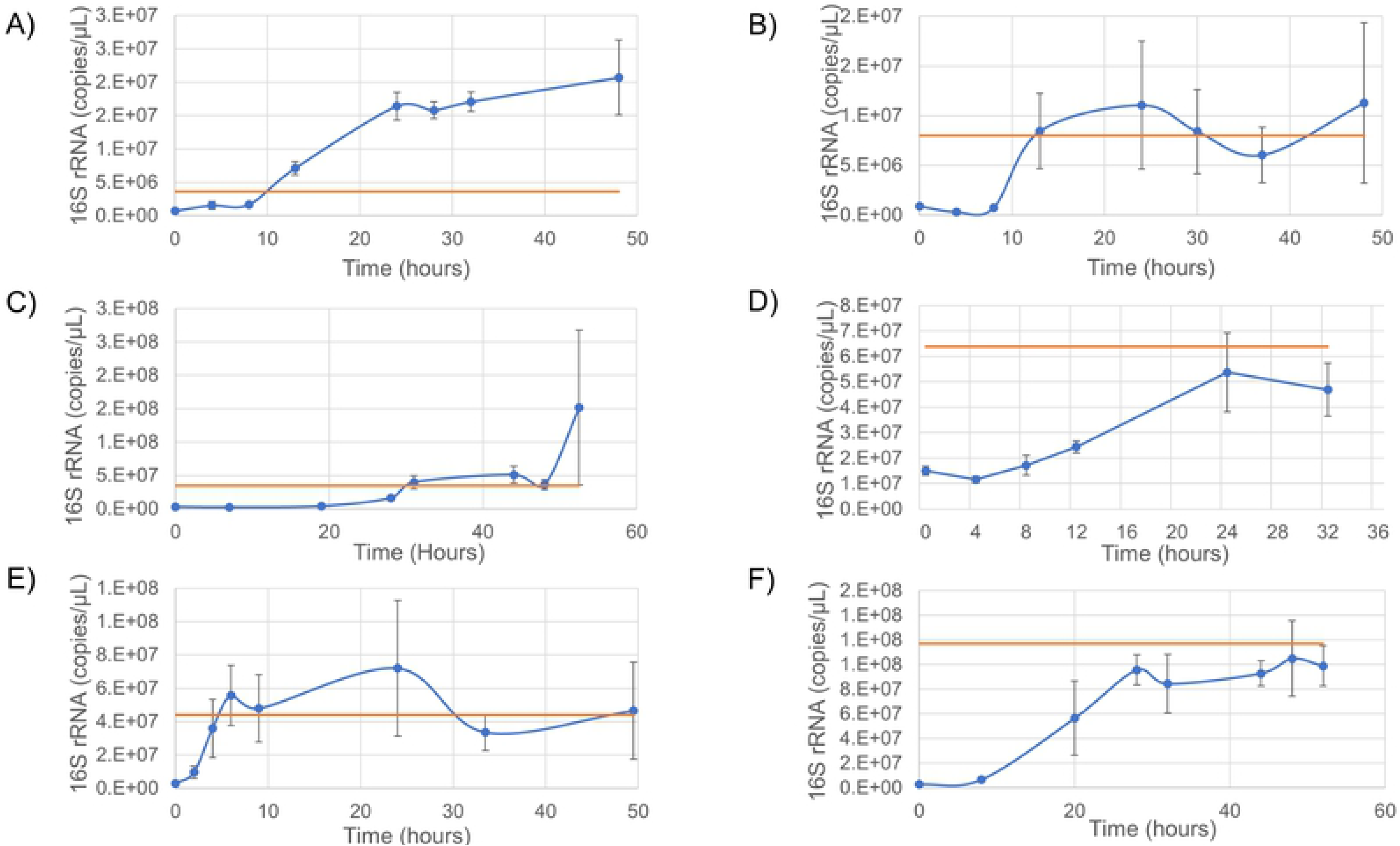
Quantitative PCR results targeting 16S rRNA of a time course for each species. 16S rRNA copies per microliter as detected by qPCR over a time course of A) *A. woodii,* B) *B. subtilis,* C) *D. vulgaris,* D) *G. subterraneus,* E) *P. putida* and F) *T. aromatica*. Blue lines are the species growth curves (n=6, three biological each with two technical replicates), orange line is the inoculant value.

Cell count equivalents for 16S qPCR readings are reported in Table 5 and were calculated using the unique 16S rRNA gene copy numbers (per cell) as reported in Table 1.

## Discussion

First let’s take the opportunity to do a ‘mini’ review of cell quantitation (count) methods for bacterial growth. In laboratory-based research, it is taken for granted the simplicity of assessing microbial growth. This is typically performed using direct counting methods such as colony forming units (CFU) plated on appropriate agar, where after each incubation each cell can be visualized as a unique colony on the agar. Other metabolic or growth specific methods exist such as dilution series (colloquially known as “bug bottles”) [35] and bacteriological activity reaction test (BART) bottles [5], both of which produce a colour change resulting from specific microbial growth and metabolism where the time until detectable change (i.e. a change in the colour of the media) is used to approximate the initial microbiological cell count. These are commercially available for sulfate reducing bacteria, acid producing bacteria, iron reducing bacteria and others. All these methods are dependent on the culturability of the microbes in question and are not applicable when the species in question cannot be cultured easily or conveniently. Due to the nature of these techniques and their estimation of culturable microorganisms and not quantitative enumeration, these techniques are more commonly used in industry where exact cell counts are not required as opposed to scientific endeavours which typically require more precise counts.

During planktonic growth, optical density (OD) is frequently used as a rapid monitoring method where the turbidity of the media is correlated to the cell density. If a sample or culture has high turbidity, the effect of light scattering by the cells is diminished and the measured OD becomes too high to provide a linear application of the Beer-Lambert law. Studies have shown that OD measurements to assess cell counts are highly dependent on the spectrophotometer, wavelength, media type, growth stage, cell morphology and the presence and concentration of secreted compounds, and thus OD measurements should be taken as a proxy and not as concrete correlations [36]. Klett units are a similar means of determining cell concentrations using turbidity, where the turbidity of a liquid culture has been correlated to a colony forming unit value, and this is commonly done on a per strain basis via wavelength filters as part of this older tool [37,38]. Direct comparison of OD for different species is difficult due to changes in turbidity resulting from cell shape such as rod compared to coccoid and cell agglomeration to flocculation issues and thus not individual cell density which can lead to incorrect assumptions at the same OD. This will not pose an issue while studying a pure culture of known shape as comparisons are direct. However, for cross comparisons or for mixed cultures this becomes a significant issue and lowers the utility and accuracy of OD as a tool for growth measurement.

Moving away from the culture-dependant methods, dyes and stains can be used to visualize and semi-quantitatively measure growth. Crystal violet is used as a biofilm assessment assay [39] but Safranin has been found to be more reproducible [40]. These stains do not provide true cell counts but produce quantifiable biomass measurements, which depending on the research question posed is sufficient to monitor microbial growth.

Other, less direct methods exist which can approximate cell counts through quantification of other components such as key metabolites including ATP or major biochemical compositions such as protein concentration, DNA concentration, or lipid composition. Each of these techniques relies on an average quantity of the target molecule being present in each cell and have similar trends and limitations as DNA, where fluctuating concentrations may be a reflection of a specific stage of cellular division, but largely the absolutely concentrations are maintained at a steady state [41–43]. Lipids can also be used to determine growth rates by tracking the incorporation of heavy water into lipids using gas chromatography [44]. In the case of ATP, assays use an average quantity of ATP per cell based on *E. coli* where one *E. coli* cell contains approximately 1 femtogram of ATP [45], while other studies have shown ATP concentrations are stable throughout all growth rates, although exact ATP concentrations per cell were not calculated [46]. These assumptions do not take into account periods of external stress (e.g. biocide exposure) or temperature increases which may increase the intracellular ATP concentrations in response [47]. An added benefit of these methods when considering non-defined environmental samples is that they are only present in biochemically active cells, and free molecules do not survive for long outside of a living cell. As such, in environmental samples these lines of evidence can be reasonably used to estimate total biomass without having to consider the types or diversity of species present.

More advanced methods of quantifying the number of organisms present in complex samples have arisen from molecular methods such as quantifying specific gene copy numbers including 16S rRNA genes, housekeeping genes other species-specific marker genes. Many databanks exist which have developed different 16S rRNA primer sets, each with their own biases towards detecting or omitting certain microbial clades and are reviewed elsewhere [48–50]. Primer sets are often chosen if there is an indication as to the types of organisms expected to be present (e.g. a predisposition towards methanogenic microorganisms). These primers can be used in quantitative PCR (qPCR) for total cell counts, or in sequencing to acquire relative abundance values.

Further expanding on the applications of qPCR is the customization of the primers used. While 16S rRNA is a universal gene target and been used dating back to 1999 [51], other housekeeping genes specific to a species or interest may be used to get cell counts of targeted populations. For example, the use of primers targeting dissimilatory sulfate reductase, *dsrA,* to monitor sulfate reducing organisms [52] or nitrite reductase, *nirS*, to quantify *Pseudomonas stutzeri* [53], aromatic oxygenases [54]and hydrocarbon hydroxylases [55]. An important consideration when using genes as a cell count proxy is the copy number of the gene in each cell. For 16S rRNA specifically, bacteria can have anywhere from a single copy up to 15, with an average of 3.82 ± 2.61 [56].

### Comparison of methods

To compare how the different methods agree with each other, scatter plots were used and linear correlation values were calculated (supplementary Figures 1-6). The R^2^ values were calculated for the full datasets of each monitoring method for each species and are reported in Table 4. From these values we can see that the different methods have stronger correlations within certain species, such as *A. woodii,* which has strong correlation between all methods (R^2^ = 0.85 – 0.99), while *B. subtilis* has poor correlation across almost all methods where R^2^ values range from 0.41 to 0.79 (with the exception of DNA vs. 16S rRNA, R^2^ = 0.96). Other species show a strong correlation between some methods such as OD_600_ vs. DNA (*G. subterraneus,* R^2^ = 0.98) but poor correlation with others (OD_600_ vs. ATP, R^2^ =0.06, and ATP vs. DNA R^2^ = 0.02).

**Table 4.** Linear correlation* values determined from scatter plots of each growth monitoring technique.

| Species | OD<sub>600</sub> vs<br>ATP | OD<sub>600</sub> vs<br>DNA | OD<sub>600</sub> vs<br>16S | ATP (ME/mL)<br>vs DNA | ATP (ME/mL)<br>vs 16S | DNA vs<br>16S |
| --- | --- | --- | --- | --- | --- | --- |
| *A. woodii* | 0.94 | 0.93 | 0.96 | 0.85 | 0.87 | 0.99 |
| *B. subtilis* | 0.79 | 0.71 | 0.74 | 0.41 | 0.44 | 0.96 |
| *D. vulgaris* | 0.43 | 0.87 | 0.55 | 0.55 | 0.25 | 0.38 |
| *G. subterraneus* | 0.06 | 0.98 | 0.97 | 0.02 | 0.02 | 1.00 |
| *P. putida* | 0.96 | 0.99 | 0.51 | 0.97 | 0.54 | 0.59 |
| *T. aromatica* | 0.94 | 0.81 | 0.90 | 0.79 | 0.84 | 0.97 |
| Average | 0.69 | 0.88 | 0.77 | 0.60 | 0.49 | 0.82 |

\*Reported as the $R^2$ value from XY scatter plots of each data set against each other.

After converting to microbial equivalents (ME), the ATP measurements follow very similar trends as the OD_600_ readings (average R^2^ = 0.69, Table 4). The most marked differences in the ATP compared to the OD_600_ graphs occurs in *D. vulgaris* (R^2^ = 0.43) and *G. subterraneus* (R^2^ = 0.06) where the shapes of the curves are significantly different owing to the peak in ATP occurring before the stationary phase. Comparing the OD_600_ and ATP values for *A. woodii* (R^2^ = 0.94), *B. subtilis* (R^2^ = 0.79), *P. putida* (R^2^ = 0.96) and *T. aromatica* (R^2^ = 0.94), we see a strong correlation with slight variations and more variability in the ATP values than seen in OD_600_. These data agree with the observation that the amount of ATP is consistent at all stages of growth curve [46], while the *D. vulgaris* and *G. subterraneus* datasets disrupt the expected sigmoidal growth curve and shows ATP concentrations are highest during mid log phase, indicating this isn’t a universal rule. This is further exemplified in the *G. subterraneus* dataset, whose ATP values were markedly different from other approaches, and correspondingly the linear correlations with ATP are all very low (OD_600_ vs. ATP R^2^ = 0.06; ATP vs. DNA R^2^ = 0.02; and ATP vs. 16S R^2^ = 0.02).

The trends in the DNA concentrations closely follow the OD_600_ trends (average R^2^ = 0.88, Table 4), but have more variance between replicates, possibly owing to the DNA extraction and recovery procedure. As with the DNA concentrations, the trends of the 16S rRNA copy numbers closely followed the OD_600_ readings (average R^2^ = 0.77) but have a lower linear correlation due to the increased variability between 16S rRNA replicates. Unsurprisingly, DNA vs. 16S rRNA had a strong correlation (average R^2^ = 0.82), but the strongest correlation between any two methods is the OD_600_ and DNA (average R^2^ = 0.88). The DNA vs. 16S correlation value is dropped by the high variability of the 16S rRNA copies for the final time point of *D. vulgaris*, but with this final time point removed the average R^2^ value improves to 0.91 (data not shown). The *P. putida* 16S rRNA vs. DNA correlation is also poor but this is again owing to the variability of the replicates for 16S rRNA during the stationary phase. ATP vs. 16S rRNA methods had the lowest correlation on average between the six pure cultures with an average R^2^ value of 0.49 (Table 4), which was mirrored in the correlation between DNA vs. ATP (average R^2^ = 0.60, Table 4).

From these comparisons, we can see that both ATP vs. 16S rRNA and ATP vs. DNA concentrations have poor correlation with each other. However, the R^2^ values of these averages is skewed downwards as a result of the extremely poor values of *G. subterraneus.* With those values removed, the R^2^ values improve to 0.71 for ATP vs. DNA and 0.59 for ATP vs. 16S rRNA. It is tempting to consider the R^2^ values with the omissions of the *G. subterraneus*, however they provide a realistic comparison of the diversity of values one might expect in an environmental sample, even in such a small pool as the six species chosen here.

### Direct comparison of cell counts

For direct comparisons, we shall look at three time points for each species, chosen to reflect the initial time point (T = 0 hours), mid-log phase (variable) and stationary phase (variable) for each species. For easier direct comparison, all testing methods have been calculated into their unique cells mL^−1^ (Table 5). CFU calculations from all four monitoring methods showed variability of a magnitude lower than the reading, however and on occasion the same order of magnitude or in the case of OD_600_ two orders below. Comparing across the ATP, DNA and 16S rRNA measurements, the ATP microbial equivalents are an order of magnitude or two below calculated DNA and 16S rRNA values. Looking at a single measurement type of a single species, the change between the initial time point to the stationary phase is typically an order of magnitude regardless of the monitoring method. *B. subtilis* had the largest discrepancies between DNA and 16S rRNA calculated cell counts, varying by an order of magnitude. The lowest calculated cell count from all species along all time points is the microbial equivalents of *B. subtilis* at T = 0 hours, which was three to four orders of magnitude below other methods but the discrepancy was closed by mid log phase and was similar to the calculated values of the other methods and species (Table 5). Stationary phases for DNA and 16S rRNA plateaued primarily at 10^9-10^ cells mL^−1^ (*B. subtilis* 16S rRNA being the exception at 8.4 × 10^8^ cells/mL). ATP and OD_600_ values plateaued at 10^7-8^ CFU/mL with the single exception of *A. woodii* ATP value (1.45 × 10^9^ ME/mL). This indicates that following these calculations, DNA and 16S rRNA calculations will likely overestimate cell counts compared to OD_600_ and ATP.

## Conclusion

This work set out to compare monitoring techniques readily employed in the field and compare them to methods best suited for lab cultures. OD_600_ is poorly suited for field samples due to the requirement for a liquid medium and the presence of non-biological materials frequently present in environmental samples which will artificially increase OD_600_ values. This is illustrated by the *D. vulgaris* dataset, where the scattering of light is increased due to the precipitation of iron sulfide resulting from sulfide production by *D. vulgaris*. Even without the presence of a precipitate in the media, direct OD_600_ comparisons have little value such as with *G. subterraneus,* where OD_600_ values peaked at 0.317 while the other species reached readings of 1.001 and 1.296 in the stationary phase (*P. putida* and *A. woodii* respectively). As a result, OD_600_ is only suited for rapid monitoring of a pure culture and has no value for cross comparisons.

As shown in Table 5, the trends are consistent but the actual converted cell counts can vary on the order of magnitudes. This indicates that while any single method can be used with reasonable confidence to assess the microbial cell density in a particular system or environment, comparing multiple methods will lead to false assumptions regarding changes in cell concentrations.

Due to the need of including additional values for converting ATP, 16S rRNA copy numbers and DNA into cell count equivalents, it is more reasonable to leave these readings as their true output (i.e. pg ATP, copy number 16S rRNA and μg DNA respectively) rather than adding converting values using equations containing general assumptions, which certainly can skew output values, especially in mixed environmental samples where the factors (e.g. DNA amount per cell) may vary between species.

This work shows that using DNA concentrations as a proxy for cell counts could be considered the best universal indicator for microbial cells numbers. It carries a strong correlation to OD_600_ values in pure cultures in liquid media, is not as susceptible to large variation between replicates as 16S rRNA qPCR and theoretically isn’t overly susceptible to calculation bias as the range of DNA concentrations within a single cell is 1.6-2.4 fg [33]. Using DNA as a cell count proxy has the only reproducible DNA recovery as a potential issue, where Gram positive cells may not lyse as easily as Gram negative cells in which case DNA concentrations may underestimate the total cell counts.

While this work focused on pure cultures of diverse environmental strains, we believe these results can be extrapolated to mixed species and field samples with highly diverse microbial populations. The simplest and most impactful conclusion from this is that there is no true, or best method for monitoring microbial growth, rather being consistent with a monitoring technique is the most important factor and to understand the used approaches limitations as illustrated here.

